# Phenotypic diversity and shared genomic determinants among isolates causing a large incidence of disseminated gonococcal infections in Canada

**DOI:** 10.1101/2024.09.08.611882

**Authors:** Gursonika Binepal, Emil Jurga, Duncan Carruthers-Lay, Sören Krüger, Sandra Zittermann, Jessica Minion, Mathew Diggle, David C. Alexander, Irene Martin, Vanessa Allen, John Parkinson, Scott D. Gray-Owen

**Affiliations:** Department of Molecular Genetics, University of Toronto, Toronto, Ontario, Canada; Program in Molecular Medicine, Hospital for Sick Children Research Institute; Department of Biochemistry, University of Toronto, Toronto, Ontario, Canada; Public Health Ontario, Toronto, Ontario, Canada; Department of Pathology and Laboratory Medicine, University of Saskatchewan, 1440 14th Avenue, Regina, SK, Canada, Roy Romanow Provincial Laboratory, (formerly Saskatchewan Disease Control Laboratory) 5 Research Drive, Regina, SK, Canada; Alberta Precision Laboratories - Public Health Laboratory (ProvLab), Edmonton, AB, Canada; Cadham Provincial Laboratory, Diagnostic Services, Shared Health, Winnipeg, Manitoba, Canada; National Microbiology Laboratory, Public Health Agency of Canada, Canadian Science Centre for Human and Animal Health, Winnipeg, Canada; Department of Respiratory Medicine, Leipzig University, Leipzig, Germany, Department of Intensive Care Medicine, University of Aachen, Aachen, Germany

**Keywords:** Neisseria gonorrhoeae, disseminated gonococcal infection (DGI), gonorrhea, sexually transmitted infection (STI), phenotypic analysis, genomic analysis

## Abstract

The incidence of disseminated gonococcal infection (DGI) has remained low since the advent of antibiotics, however recent surge in DGI have inexplicably emerged within several regions during the past decade. In an effort to understand whether *Neisseria gonorrhoeae* that cause disseminated disease can be differentiated from non-invasive strains, we have performed a phenotypic and genotypic analysis on a selection of isolates obtained from invasive and uncomplicated infections in Canada. Phenotypic analysis of a matched subset of 19 isolates obtained since 2013 found that these varied in their capacity to aggregate in suspension and in their association with serum complement proteins, however these interactions did not discriminate between the invasive and mucosal isolates. Sequence typing of 360 Canadian isolates revealed that two *porB* alleles are significantly associated with the DGI strains, one of these being present throughout the past decade whereas the other became associated more recently. A PopNet-based population dynamics analysis, which instead establishes relationships based upon variance among discrete chromosomal segments, found that DGI isolates were restricted in their phylogenetic distribution. While this implies a genetically-linked potential to cause invasive disease, it cannot distinguish between an inherent difference in the phenotype of these populations or the horizontal exchange of some virulence factor among closely related strains. Regardless, a large number of genetic determinants are enriched in the DGI strains, making these enticing candidates for future work to understand how they might either promote the gonococcal capacity to cause systemic infection or reduce the presentation of clinical symptoms from localized infection so that it remains untreated.

**AUTHOR SUMMARY:** *Neisseria gonorrhoeae* is a sexually transmitted bacteria that causes over 82 million cases of gonorrhea each year. With its re-emergence, rising incidence rates and the prevalence of multidrug resistant strains increasing, the bacteria is considered to be a high burden threat to global public health. While disseminated gonococcal disease arising from untreated infections are uncommon, there have recently been regions with high incidence invasive disease. Here, we take advantage of the active ongoing collection of gonococcal isolates in Canada to perform a combined phenotypic and genotypic analysis that aims to understand whether certain strains are more often linked to invasive infections. We found that all disseminated isolates bound to the complement regulatory factors factor H and/or C4 binding protein to facilitate their resistance to the bactericidal activity of serum, but this was not sufficient to explain the heightened instance of invasive disease. While classical genome-based phylogenetic analysis displayed little association between invasive strains, PopNet-based analysis revealed that invasive isolates fell within defined sub-populations and indicated variant alleles enriched among the disseminating bacteria, and a broader pan-genome approach revealed genes more likely to be present in invasive strains. Our study thereby provides support for a genetic contribution to the invasive potential of gonococcal isolates and provides candidate drivers of this virulent outcome.

## INTRODUCTION

*Neisseria gonorrhoeae* (*Ngo*) causes over 82 million infections per year globally [1]. This high burden of gonococcal infection and its progressive acquisition of resistance to antibiotics has created an increased urgency for understanding its epidemiology and developing new countermeasures to combat infections [1]. Symptomatic infections typically present with localized urogenital manifestations including profuse urethral or cervical discharge, though asymptomatic colonization of these or other mucosal sites is common. The highest burden of disease occurs in untreated women, where *Ngo* can spread into the upper genital tract to cause inflammation and tissue scarring, with sequelae including chronic pain, ectopic pregnancies and infertility [2]. In rare cases, disseminated gonococcal infections (DGI) can arise in both men and women, with presentations including migrating arthritis, tenosynovitis, endocarditis and/or dermatitis [3, 4]. DGI has been reported to occur in 0.5%-3% of those infected [3, 5, 6], though other studies suggest that rates are now typically below this [4], presumably due to the success in treatment preventing disease progression. Recently, unexplained clusters of high DGI incidence have been reported in Manitoba, Michigan, Georgia and California [7–10]. While the rates of gonococcal infection have increased, the proportion of cases that manifest as DGI has been significantly higher in these regions without obvious increases in other areas. Also notable is that DGI has historically been higher in women, while these more recent emergences appear to affect men and women equally, suggesting some unique characteristics of these recent cases.

Gonococcal isolates from disseminated sites have long been known to resist the bactericidal activity of normal human serum [11]. This has been attributed, at least in part, to the unique ability of gonococcal strains to variably bind complement regulatory factors that normally function to protect human cells from complement-dependent killing, including factor H (FH), C4b binding protein (C4BP) and vitronectin. Gonococcal sialylation of its lipo-oligosaccharide (LOS), neisserial surface protein A (NspA) and the porin B (PorB) variants can each, synergistically, promote FH binding to the bacterial surface, which confers resistance to the alternative (immunoglobulin-independent) complement cascade (reviewed in [12]). Certain variants of PorB also bind human C4BP, which inhibits immunoglobulin-dependent (classical) complement killing. C4BP binding is most commonly apparent with expression of a *porB1a* allele. This is not a strict association given that the interaction is evident in 80% of strains expressing PorB1a and 20% of those expressing the closely related but antigenically distinct PorB1b [12], yet the relationship is consistent with the fact that invasive isolates tend to express PorB1a [12, 13]. The diversity of mechanisms by which the gonococci can decorate their surface with complement regulatory factors to resist the bactericidal activity of serum, and the absence of any other factors known to increase the propensity of *Ngo* to cause systemic infection, makes it difficult to understand why individual strains may be more or less likely to cause DGI.

Given the genetic diversity of *Ngo*, it is notable that phylogenetic analysis of the isolates recovered from the DGI cases in Michigan [7] and Manitoba [8] indicates that they are highly related, with the latter being largely associated with *Ngo* having closely related multi-antigen sequence types (NG-MAST), while those isolated in Georgia were more genetically diverse [10]. A marked increase in DGI has also been apparent in Ontario [14], which abuts both Manitoba and Michigan, first emerging in 2014 and then remaining at elevated rates since that time. Genomic sequence analysis by Public Health Ontario (PHO) indicates that the early strains were of the NG-MAST that has been associated with most DGI in Manitoba (type 11508), suggesting that they are epidemiologically linked, however latter strains have diverged from this. Herein, we have taken advantage of the routine collection of gonococcal isolates by public health agencies in Canada to characterize the relative capacity of disseminated and non-disseminated (mucosal) isolates from 2013 through 2019 to bind complement factors and resist the bactericidal activity of human blood, and to search for genetic determinants that are uniquely shared by the diverse isolates recovered.

## RESULTS

### Serum resistance of low passage *Neisseria gonorrhoeae* isolates

Disseminated gonococcal infections are typically considered to emerge from asymptomatic and/or untreated genital infections. The propensity for certain strains to frequently be associated with invasive infections suggests that they possess some capacity that is not shared by strains that remain localized to the genital or other mucosal surfaces. To consider what might distinguish isolates that do or do not have the propensity to cause invasive disease, we selected a subset consisting of 19 clinical isolates obtained between 2013 and 2019, including 14 isolated from invasive infection and 5 isolates with similar NG-MAST type but from uncomplicated genital infection (**Table 1**). Of these, 10 were from Ontario, 6 from Manitoba, 2 from Saskatchewan and 1 from Alberta. Both alleles of porin B, the most abundant antigen in the outer membrane and key element of molecular typing schemes, are represented, with 14 PorB1a and 5 PorB1b isolates. These attributes are depicted relative to their phylogenetic relationship, based upon concatenated single nucleotide polymorphisms generated after alignment with the FA1090 genome, in **Figure 1A**.

**Table 1:** Summary of isolates used in the study.

|  |  | Number of Samples |  |  |
| --- | --- | --- | --- | --- |
|  |  | Total | Non-DGI | DGI |
| Year |  |  |  |  |
|  | 2013 | 1 | 0 | 1 |
|  | 2014 | 9 | 5 | 4 |
|  | 2015 | 135 | 131 | 4 |
|  | 2016 | 80 | 77 | 3 |
|  | 2017 | 53 | 34 | 19 |
|  | 2018 | 67 | 38 | 29 |
|  | 2019 | 15 | 1 | 14 |
| Gender |  |  |  |  |
|  | Male | 272 | 232 | 40 |
|  | Female | 81 | 49 | 32 |
|  | Unknown | 7 | 5 | 2 |
| Age |  |  |  |  |
|  | 10-19 | 26 | 19 | 7 |
|  | 20-29 | 143 | 126 | 17 |
|  | 30-39 | 92 | 73 | 19 |
|  | 40-49 | 64 | 46 | 18 |
|  | 50-59 | 14 | 10 | 4 |
|  | 60-69 | 12 | 6 | 6 |
|  | 70-79 | 2 | 1 | 1 |
|  | Unknown | 7 | 5 | 2 |
| PorB allele |  |  |  |  |
|  | porBa | 115 | 60 | 55 |
|  | porBb | 230 | 224 | 6 |
|  | Unknown | 15 | 2 | 13 |
| Province |  |  |  |  |
|  | Ontario | 294 | 254 | 40 |
|  | Manitoba | 59 | 27 | 32 |
|  | Saskatchewan | 5 | 3 | 2 |
|  | AB | 2 | 2 | 0 |

**Figure 1.**
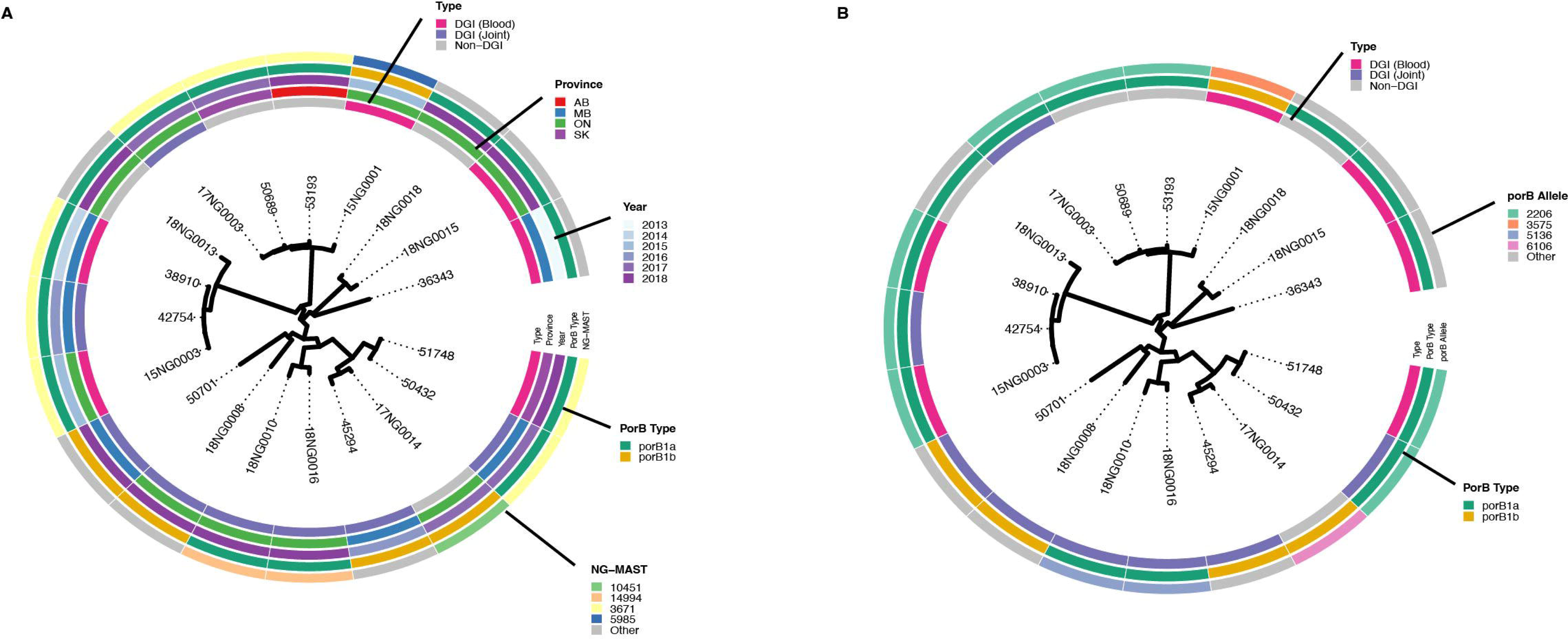
Characteristics of *N. gonorrhoeae* clinical isolates. **(A)** Maximum likelihood tree of 19 isolates selected for phenotypic analysis constructed from a concatenated set of SNPs generated from alignment to the FA1090 reference genome. Only branches with bootstrap support >0.8 are shown. Rings (outside-in) represent NG-MAST serotypes, PorB allelic variant, year of collection, province and DGI phenotype. **(B)** Outer rings (outside-in) represent *porB* allele, *porB* type and DGI phenotype

Given that resistance to the bactericidal activity of complement is presumed to be a prerequisite for systemic infection, we first undertook to assess the susceptibility of these isolates to human serum. To optimize the serum concentration for use in the assays, we selected 6 different strains (15NG0003, 17NG0014, 18NG0016, 36343, 50412 and 53193) and then exposed these to dilutions ranging from 10% to 90% human serum (HS; **S. fig. 1**). Based on colony forming count data, a concentration of 60% HS was bactericidal for 2 of 6 strains, so was selected as a basis for differentiating serum sensitivity between the isolates. Applying this to the full set of 19 clinical isolates, we found varying levels of serum resistance, either in the presence or absence of 5′-cytidinemonophospho-*N*-acetylneuraminic acid (CMP-NANA) (**Figure 2**), which some strains can use to sialylate their LOS and thereby increase their survival [12]. Specifically, three of the five isolates from uncomplicated infection were resistant in human serum, and another became resistant in the presence of CMP-NANA (4 total, 80%), while 7 of 16 DGI isolates were resistant in serum and 4 more became resistant with CMP-NANA (11 total, 69% resistance). Six of 14 PorB1a isolates were resistant to serum and an additional 4 became resistant in the presence of CMP-NANA (10 total, 66% resistance). Two of the 3 DGI isolates that express PorB1b were resistant to serum while the other was sensitive both in the presence and absence of CMP-NANA (66% resistance), and the mucosal isolate that expresses PorB1b was also sensitive in both conditions.

**Figure 2.**
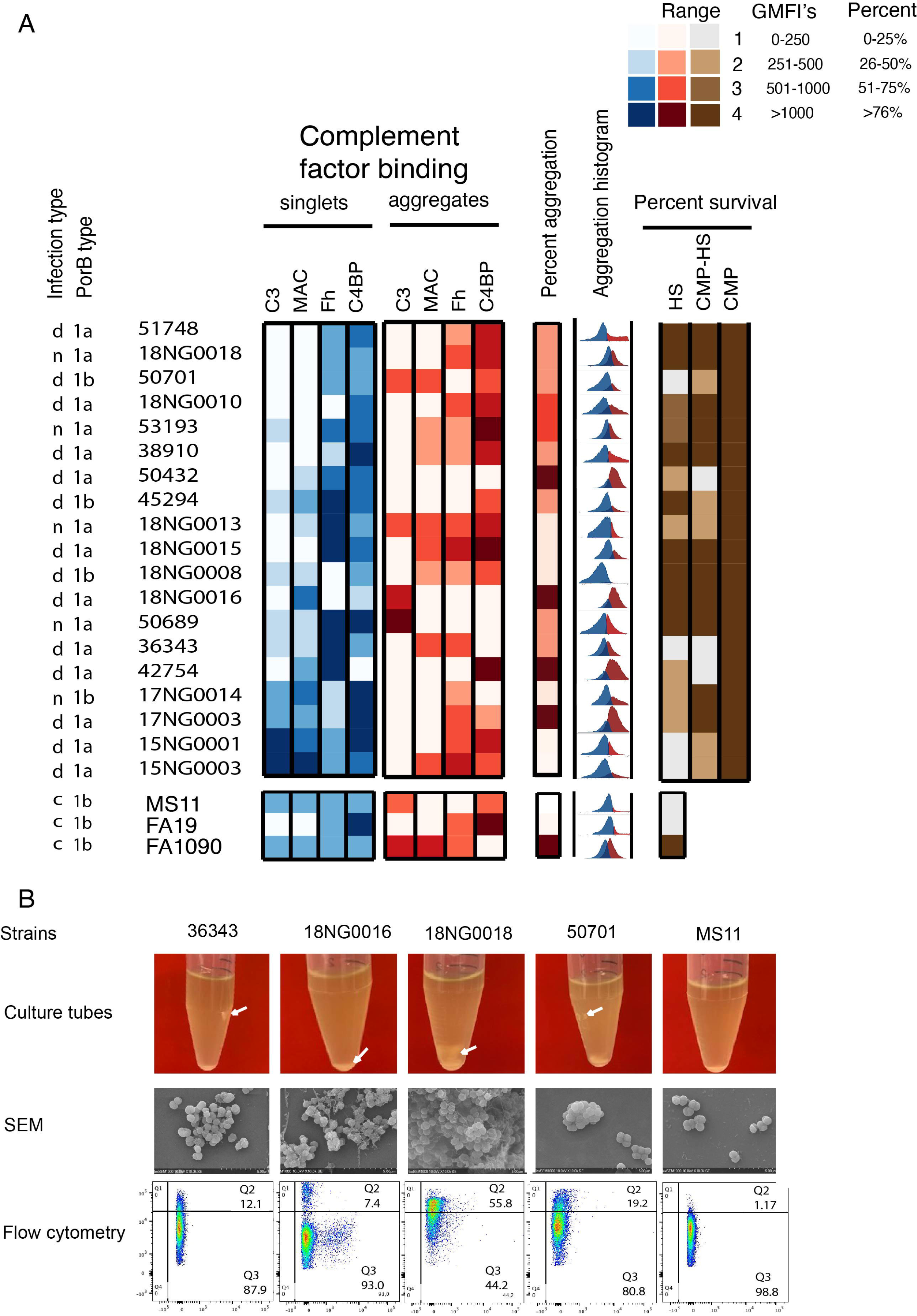
Serum resistance and complement binding assays. **(A)** PorB type and infection type (disseminated (d) or non-disseminated mucosal (n) are shown for each strain. Binding of indicated (C3, C5b-9 complex of the membrane attack complex (MAC), factor H (FH) and C4b binding protein (C4BP)) complement factors to bacterial singlets (blue) and aggregates (red) after treatment with 60% HS are shown as heat maps. The percentage of singlet versus aggregated bacteria observed in normal (no serum) media (Percent aggregation) is indicated beside flow cytometry plots (Aggregation histogram) illustrating the shift in population from singlets (blue) to aggregates (red). Serum resistance was calculated to obtain percent CFU recovery (Percent resistance) and then plotted as a heat map based upon quartiles indicated in the legend. Effect of culturing in CMP-NANA (CMP) and/or human serum (HS) on the percent survival of indicated strains is shown. Values represent normalized fluorescent values (geometric mean fluorescence intensity: 0-250, 251-500, 501-1000, >1000). **(B)** Aggregation of gonococci in normal growth media. Representative gonococcal strains were grown in BHI supplemented with IsoVitalex at 37°C shaking at 180rpm for 6-8 hours. Images depict differential bacterial settling (clumping) in the culture tubes. Scanning electron micrographs of the bacterial suspensions showing clumping. Flow cytometry dot plots gated to illustrate singlets (bottom right quadrant) and larger aggregates indicative of clumping (upper right quadrant).

Sawatzky *et al* found the *porB-2206* variant allele to be typically expressed by *Ngo* isolates associated with recent outbreak of DGI in Manitoba [8]. In our subset, 7 out of 15 invasive isolates carried the *2206* allele (**Figure 1B**), of which 5 isolates exhibited high serum resistance. Interestingly, the 2 non-invasive isolates (53193 and 50689) that carried this allele (**Figure 1B**) were also highly serum resistant. The *porB-5136* allele was observed exclusively in two invasive isolates (18NG0010 and 18NG0016) that exhibited high serum resistance, while *porB-3575* was only present in one invasive isolate.

### Complement factor binding does not differentiate invasive and genital isolates

Considering that *in vitro* serum bactericidal activity (SBA) alone cannot reveal how *Ngo* interacts with individual serum components, we next sought to quantify complement protein deposition. In addition to monitoring the two human-derived negative regulators of complement activation, C4BP and factor H, both of which facilitate *Ngo* complement resistance [12], we used bacterial binding to C3 and the presence of a C5b-9 complex-dependent neoepitope as direct measures for opsonization and membrane attack complex (MAC), respectively. During these analyses, we noted a tendency for some strains to form aggregates when suspended in human serum (**S. fig. 2B**). We have depicted this as an ‘Aggregation Histogram’, which differentiates the single cell (blue) versus aggregate (red) populations, and quantified ‘Percent Aggregation’ to reflect the propensity of strains to aggregate based upon frequency of single cells (singlets) versus aggregates **(Figure 2A)**. Since aggregation would alter surface accessibility to complement factors, we evaluated binding activities for aggregates and non-aggregates (singlets) separately in **Figure 2** (see Methods). Interestingly, all four strains that displayed the highest aggregation were invasive (denoted as ‘d’ in Infection Type), however two of these (50432 and 42754) were highly susceptible to the bactericidal activity of serum in both the absence and presence of CMP-NANA.

While individual strains displayed marked variation in their level of binding to C3 and MAC, we observed no distinction between DGI and mucosal isolates, and no clear difference between PorB1a and PorB1b-expressing strains (**Figure 2A, Supp. Figure 1A, C, D**). Most strains displayed substantial binding to FH and/or C4BP and low or no C3 or C5b-9 binding, consistent with their being highly resistant to bactericidal killing. One outlier in this regard is strain 18G0008, which showed little interaction with any complement or complement regulatory proteins and was serum resistant. Five strains were highly susceptible to killing by serum (Percent resistance) and this was not affected by pre-culturing the bacteria in CMP-NANA. Of these, strains 50701 (DGI, PorB1b), 18G0013 (non-DGI, PorB1a) and 36343 (DGI, PorB1a) displayed very low binding to C3 and MAC (though 36343 aggregates did display low MAC). Singlets of 15G0001 (DGI, PorB1a) and 15NG0003 (DGI, PorB1a) display high binding to C3 and MAC but remain viable in serum, perhaps attributable to the fact that aggregates of these strains did not display MAC assembly. Based upon this analysis, the increased propensity to cause invasive gonococcal infections is not reflected by any strict correlation with serum resistance or complement factor binding.

### DGI isolates are from diverse clonal lineages

To extend upon our phenotype analysis on the selected strain subset, we aimed to take an unbiased approach in consideration that an as yet undescribed function might contribute to the DGI phenotype. For this purpose, we took advantage of the availability of genome sequence data collected from a large number of isolates associated during surveillance for the ongoing incidences of DGI in Manitoba and Ontario. Of the 360 isolates included in these analyses, 184 were recovered from genital infections, 48 from rectal infections, 25 from eye infections, 20 from the throat, 13 have unconfirmed site of isolation and 74 reported as DGI (**Table 1** & **Supp. Table 1**). While most (272 of 360) of these isolates derived from male patients, the DGI isolates were relatively evenly divided between males (40) and females (32), reflecting that seen recently in these regions.

To assess the genetic variation of the isolates, genome sequences for each isolate were aligned to that of the reference strain, FA1090, a serum resistant strain originally isolated from the cervix of a patient with disseminated gonococcal infection [15]. Based on concatenated alignments of the single nucleotide polymorphisms (SNPs) between these sequences, we constructed a phylogenetic tree based on maximum likelihood (**Figure 3A**). While the earliest isolates (2013-2015) tend to be from Manitoba, we otherwise observe little population structure associated with geography, *porB* allele or NG-MAST, reflecting the inherent genetic diversity of gonococcal populations [16], even within this relatively restricted region. NG-MAST allele typing of the *porB* gene revealed 2 *porBa* alleles, *2206* (45/90 DGI) and *5136* (15/19 DGI), are significantly associated with DGI isolates (Fisher’s Exact Test, p < 0.00001). Allele *2206* was found in isolates across Canada (56 from MB, 27 from ON, and 7 from SK/AB) while all isolates with allele 5136 were collected in Ontario. Both alleles were represented in DGI isolates collected from blood and synovial fluid, although more isolates with 5136 were recovered from synovial fluid as opposed to blood (10 in synovial vs. 4 in blood).

**Figure 3:**
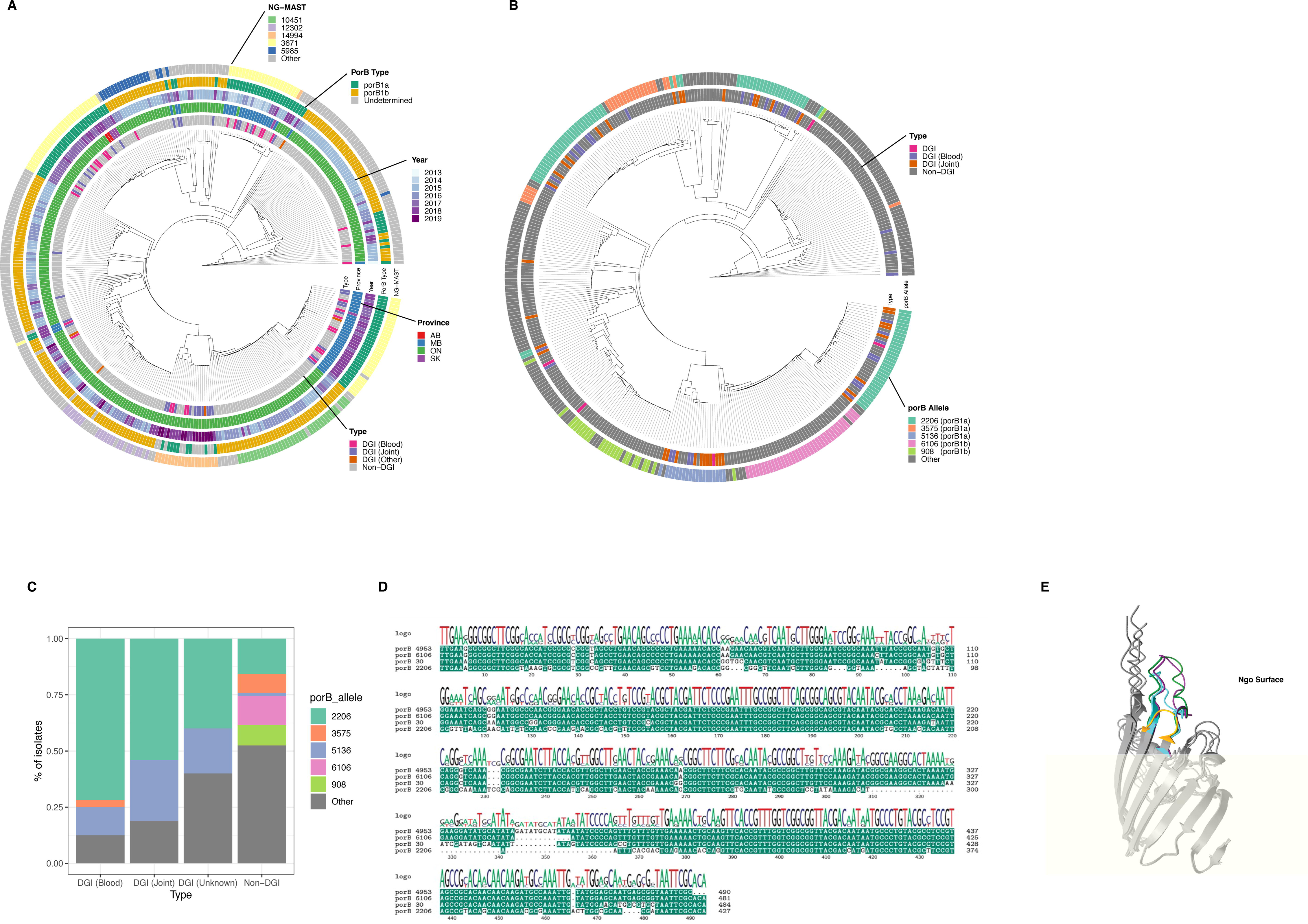
Phylogenetic relationship between 370 *N. gonorrhoeae* isolates. Maximum likelihood tree of 370 isolates constructed from a concatenated set of SNPs generated from alignment of isolate genomes against the FA1090 reference genome. Only branches with bootstrap support >0.8 are shown. **(A)** Outer rings (outside-in) represent NG-MAST serotypes, *porB* allelic variant, year of collection, province and DGI phenotype. **(B)** Outer rings (outside-in) represent *porB* allele and DGI phenotype. **(C)** Proportion of isolates carrying *porB* alleles associated with DGI phenotypes. Only the 5 most common *porB* alleles are shown. **(D)** Sequence alignment of 4 *porB* alleles; 2206 (*porBa)*, 6016 (*porBb*, non-DGI), 30 (*porBb*, DGI-associated), and 4953 (*porBb*, DGI-associated). **(E)** Structural alignment overlaying 4 *porB* alleles; 2206 (yellow), 6106 (cyan), 30 (green), and 4953 (purple). The variable loop 5 region is coloured to highlight allele-specific differences.

Of the 74 DGI isolates included in this study, 55 possessed the *porB1a* allele, which has previously been linked to the invasive phenotype. While we do not have the *porB* status for 13 DGI isolates, we note that six DGI isolates possess the *porB1b* allele, again suggesting that additional factors must be associated with the ability to cause invasive disease. In total, 13 different *porB* alleles were associated with DGI isolates **(Figure 3A-C)**. We identified 5 DGI-associated *porB1b* type alleles (3575, 6435, 30, 4953, 6892), and compared their sequences to the most abundant non-DGI associated *porB1b* allele, 6106, focusing on the variable loop 5. Alleles 30, 3575, and 6435 were highly similar, with point mutations observed in positions 330-336. We also observed a 9 base pair insertion at position 337 in allele 4953. **(Figure 3D)**. Allele 6892 was identical to allele 6106 except for a point mutation (A to G) at position 331, and was therefore not shown. Allele 30 was chosen to represent alleles 30, 3575, and 6435. In considering how the DGI-associated PorB1b proteins compared to DGI-associated PorB1a, we looked at the structural properties of these variants **(Figure 3E)**, with the grey representing mostly static regions and the variable loop 5 region is coloured with *porB1a* alleles *2206* (yellow), non-DIG associated *porB1b* allele 6106 (blue), and the DGI-associated *porB1b* alleles 30 (green) and *4953* (purple). While the localization of these sequence changes, being situated within the region deleted in *porB1a* alleles, may indicate some alterations in PorB function, it remains unknown whether its binding and/or other properties would be affected.

### PopNet analysis identifies subpopulations associated with the DGI phenotype

Based upon *porB* typing and genome-wide phylogenetic analyses shown above, it is clear that the DGI phenotype is widely dispersed across the population rather than being a single clonal lineage. Together these findings are consistent with a panmictic population structure for *Ngo* [17], but provided little insight into the relationship between DGI isolates. Given the limitations of traditional phylogenetic methods to effectively account for the impact of horizontal gene transfer (HGT) in rapidly evolving populations, we next took advantage of PopNet [18] to better understand population dynamics. PopNet divides the chromosome of each isolate into discrete segments, typically 5000 base pairs in length, and uses their pattern of single nucleotide polymorphisms (SNPs) to assess the similarity of each segment to equivalent segments in other isolates; this allows the identification of homologous regions across populations. The network visualization generated by this approach reveals relationships by color-coding each segment based upon the sub-populations that share it. Applying PopNet to the 360 isolates, we identified 8 distinct sub-populations, assigned A-H. As evident in **Figure 4A**, populations A (red), C (green), and F (yellow) cluster together, while the others are more distinct. The three central co-clustered sub-populations share patterns of variation across multiple regions of the chromosome, as indicated by their circular genomes each being interrupted by fragments of the other two colors. Isolates representing B, D, G, and H each cluster into their own populations. These are bridged to the other populations via edges (lines) that illustrate linkage to isolates from population A, perhaps indicating descendants of ancestors to these emergent sub-populations (**Figure 4A - inset**). The relative lack of colored segments in sub-populations B (blue) and H (pink) reinforces the relative isolation of these two sub-populations since it indicates a lack of genetic exchange with other groups. Isolates from sub-population E (orange) do not resolve edges to other populations at the threshold (>15% similarity) shown, indicating that it possesses the lowest genetic similarity to the other sub-populations, consistent with the fact that the genome maps are uniformly colored.

**Figure 4.**
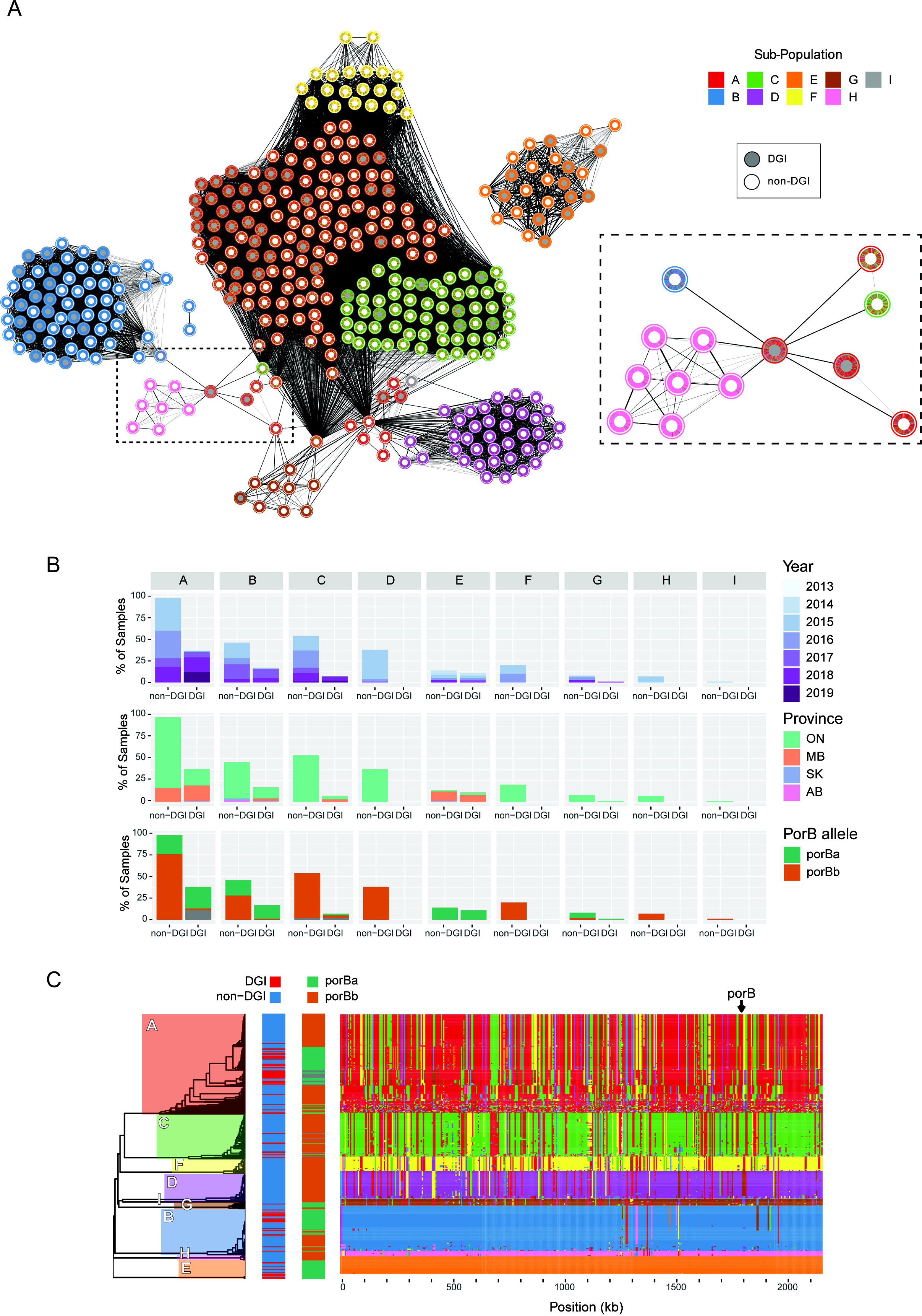
Detailed analysis of sub-populations defined by PopNet. **(A)** PopNet representation of isolates. In this network, nodes represent individual isolates and edge (line) thickness represents the percent similarity, in terms of shared SNPs. Only edges representing >15% similarity (defined as the percentage of segments that co-cluster) were used to construct the network. Nodes (isolates) are represented by annuli where colored segments represent regions of the chromosome which share ancestry across sub-populations. Filled (grey) or open (white) shading within the annuli indicate whether the isolate was associated with DGI or not. Inset, zoom in of the indicated (dashed line) set of isolates showing how one isolate links between multiple (red, blue, pink and green) sub-populations, suggesting that the isolate may be a descendant of an ancestral isolate responsible for emergence of these sub-populations. **(B)** Proportional analysis of the attributes associated with each sub-population, showing year of collection, province of isolation and PorB allele type. Percent of sample shown is calculated relative to the largest group of strains (sub-population A non-DGI). **(C)** Linear chromosome painting-based representation of PopNet-defined sub-populations that were depicted in (A). Each row represents the genome of a single isolate, clustered into their respective sub-populations. Each column represents a 5000 bp section. Section colors indicate the population to which that section shares ancestry.

Sub-populations A and C both contain isolates associated with DGI (grey-filled centers. Despite their close evolutionary relationship with sub-population F, no isolates in this latter group were associated with DGI. On the contrary, despite its apparent genetic isolation, sub-population E was found to be significantly enriched in DGI isolates relative to the overall population distribution (p=0.04, **Table 2**). This distribution confirms that the DGI phenotype has not evolved in a single clonal lineage, but instead has emerged separately in multiple lineages and/or the genetic locus responsible for this phenotype has been transmitted between the populations. It is notable that population E tends to have been isolated in the earlier years and consists primarily of samples from Manitoba (19 of 25 isolates; **Figure 4B**), making it interesting to consider that these contributed genetic material that led to the emergence of the broader DGI outbreak in Canada. The DGI isolates exhibited a greater proportion of *porB1a* allele than the non-DGI isolates, consistent with previous studies [8, 10, 12, 19]. Conversely, sub-populations D, F, H and I all lack DGI isolates and represent groups restricted to Ontario, largely isolated in 2015 and restricted to the *porB1b* allele. Beyond this, each sub-population with DGI isolates (A, B, C, E and G) includes isolates obtained over multiple years from several provinces, so they are not strictly distributed by region or date.

**Table 2:** Incidence of DGI-isolates across PopNet-defined sub-populations. This table presents a statistical analysis for the frequency of DGI isolates across PopNet-defined sub-populations that contain them. Significant enrichment for DGI-isolates was performed using Fisher’s test.

| Sub-population | Total Isolates | DGI-Isolates | p-value | Adjusted p-value | Odds Ratio |
| --- | --- | --- | --- | --- | --- |
| A | 98 | 38 | 0.09 | 0.19 | 0.67 |
| B | 63 | 17 | 0.25 | 0.31 | 0.7 |
| C | 61 | 7 | 0.11 | 0.19 | 1.99 |
| **E** | **25** | **11** | **0.01** | **0.04** | **0.33** |
| G | 9 | 1 | 0.69 | 0.69 | 2.07 |

### Genetic associations among the DGI strains

To better focus in on genetic elements with the potential to contribute to the DGI phenotype, we further examined patterns of shared ancestry across chromosomal segments (**Figure 4C**). Consistent with the network figure (**Figure 4A**), sub-population E is largely homogenous with limited shared ancestry to isolates from other sub-populations, indicating genetic isolation from other sub-populations. In contrast, sub-populations A, C, D and F share many regions with common ancestry, with A and C sharing the greatest number of regions; this again reflects the network visualization and indicates relatively frequent exchange of genetic material. Sub-population B exhibits a limited number of shared regions, indicating relatively little exchange of genetic material with other isolates. Sub-populations D, F and H, which did not contain any DGI isolates, possess few segments shared with A and C. Perhaps the most striking feature apparent is that sub-population B has a high proportion of DGI isolates and yet shares relatively few segments with the others that contain DGI strains (A, C and E); these shared regions appear interesting to explore for a potential contribution to the invasive potential.

Having identified regions of common ancestry, we next sought to identify sequences that were unique to the invasive isolates. We identified 2,657 non-synonymous SNPs associated with the DGI phenotype (p <0.20, Fisher’s exact test with Benjamini and Hochberg (BH) false discovery rate correction; **Supp. Table 2**). These variants occurred within 975 genes. Pathway enrichment analysis of the 2,657 variant SNPs revealed that they fell within 45 Gene Ontology categories (**Supp. Fig. 3**), including 12 associated with tRNA activity (e.g., methionyl−tRNA aminoacylation and ligase activity, tRNA pseudouridine synthesis). Also enriched are multiple pathways associated with the cell surface (e.g., polysaccharide biosynthesis, regulation of cell shape, cell wall organization and lipid A biosynthesis), highlighting the variability of surface proteins among strains and, possibly, between strains possessing DGI potential.

### Non-core genes may also contribute to the DGI phenotype

The analysis of genomic variants described above only considered genes that were shared with the FA1090 reference strain. However, while FA1090 was originally isolated from a patient that had disseminated infection, the repeated passaging of the prototype strain might have resulted in loss of genetic determinants not required for *in vitro* growth. FA1090 also lacks several mobile elements commonly found in gonococcal strains, including the conjugative plasmid, β-lactamase plasmid and the gonococcal genetic island, which could play a role in infection and dissemination. To account for this, we next examined whether any genes not encoded by FA1090 were enriched in the low passage DGI isolates. For each isolate, we therefore used SPAdes (v.3.15; [20]) to assemble reads that did not align to the FA1090 genome into contiguous sequences (contigs), discarding any contigs with less than 3X coverage. From the 48,788 contigs generated from all isolates, GeneMark (v 1.14_1.25_lic; [21]) annotated 91,596 putative genes. Of these, 38,379 were annotated to known orthologs through mapping to the EggNog database of orthologs (v. 5.0; [22]). These comprised 5,431 unique orthologs, of which 68% were identified in only a single isolate. Focusing on the 259 unique orthologs present in at least 5 isolates, we identified 63 non-FA1090 genes that are enriched in DGI isolates relative to non-DGI isolates (**Table 3**). Notable among them are genes related to bacterial toxin/antitoxin systems (*e.g.* putative antitoxin of a bacterial toxin-antitoxin system YdaS/YdaT, and ParE toxin of type II toxin-antitoxin system, *parDE*), as well as genes involved in conjugation and DNA transfer (*e.g.* TrbL/VirB6 plasmid conjugal transfer protein, bacterial conjugation TrbI-like protein, P-type DNA transfer ATPase VirB11). While the former may indicate some differential response to stress, the latter set of genes may suggest that DGI isolates of *Ngo* are more capable of participating in HGT.

**Table 3:**
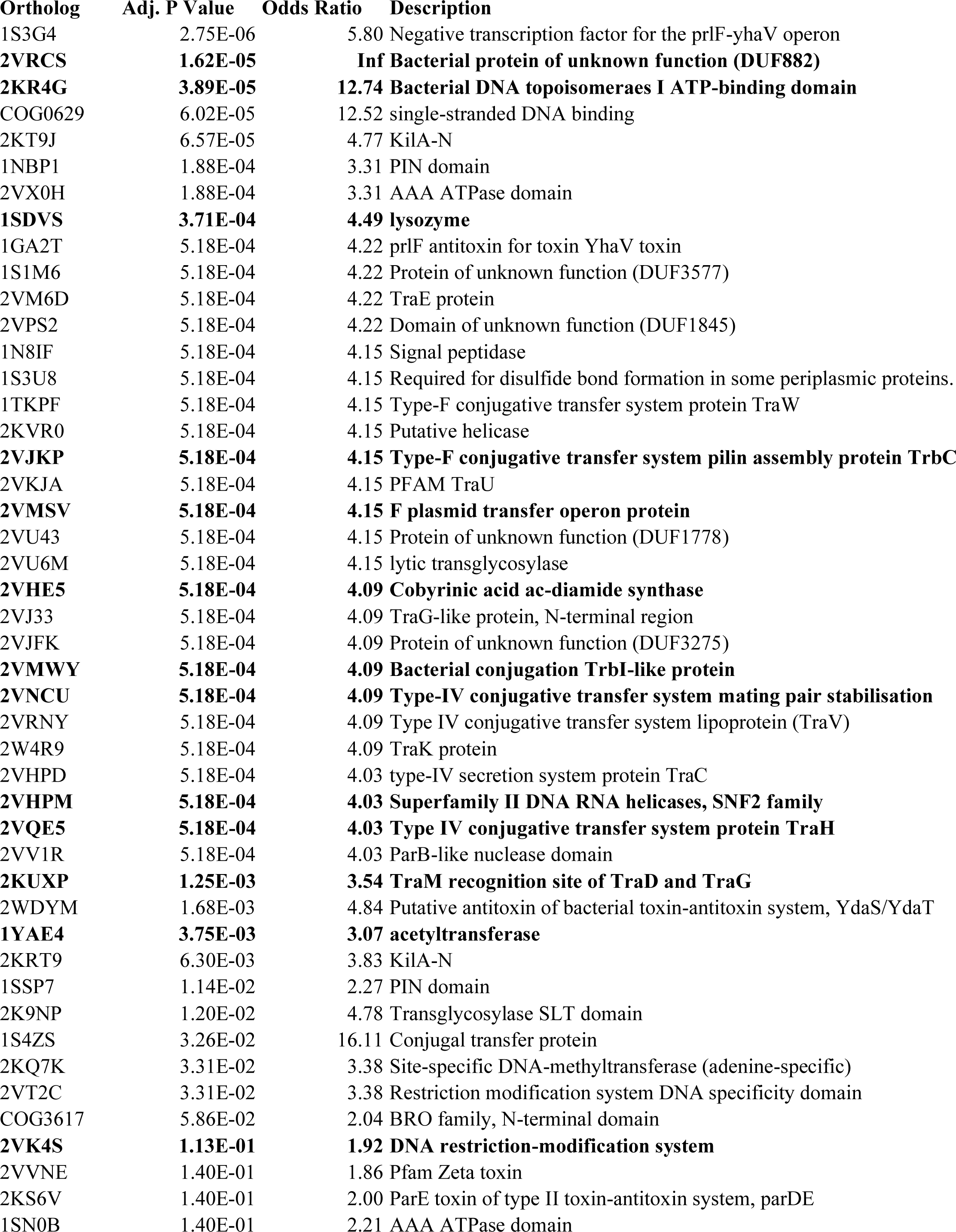
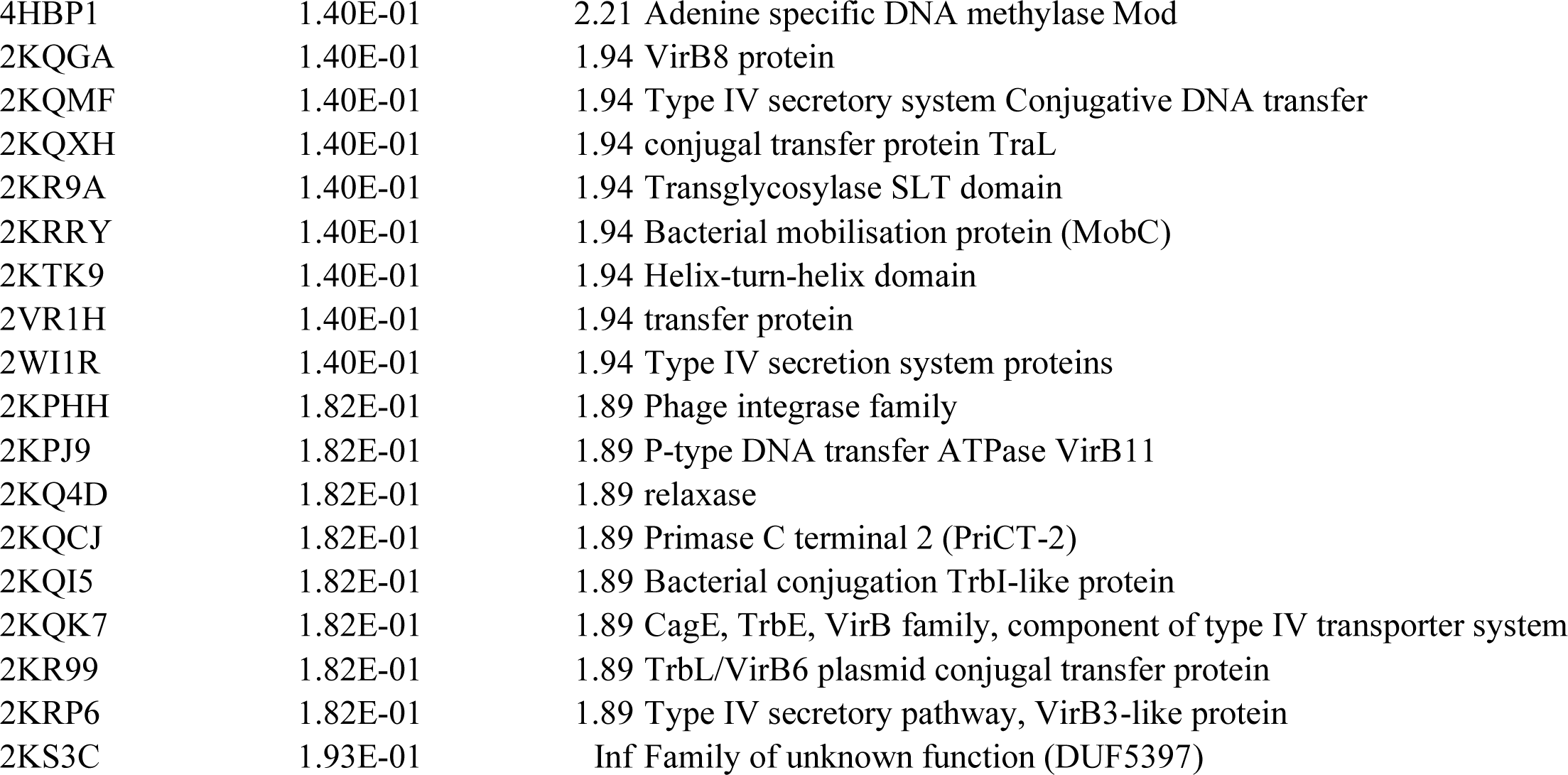
Non-FA1090 genes that were more likely to be found in DGI isolates then non-DGI isolates. *ID refers to the ortholog identification number from the EggNog 5.0 database. Bolded genes are present in the Gonococcal Genomic Island.

Among the set of DGI enriched-genes, and somewhat provocatively, the highest odds ratios are associated with two bacterial protein of uncharacterized function (DUF882(has a predicted capboxypeptidase domain) and DUF5397) (**Table 3**); four other significant DGI-associated proteins also had no known function. Beyond this, 2 of the top 3 associated proteins are related to nuclease function, including a transcription factor proposed to negatively regulate the type II toxin-antitoxin system comprised of PrlF and the YhaV translation-dependent ribonuclease (ortholog 1S3G4; the PrlF antitoxin (1GA2T) was also highly associated) [23] and a type I topoisomerase (ortholog 2KR4G), while the 4^th^ and 5^th^ are a single-stranded DNA binding protein (COG0629) and DNA binding PIN domain (1NBP1).

A large number of DGI-enriched genes were related to Type IV secretion systems (T4SS). Many strains of *N. gonorrhoeae* encode a 57-kb gonococcal genetic island (GGI) containing up to 63 genes, which encodes a T4SS responsible for the extracellular release of linear single-stranded DNA [24]. Since our reference strain (FA1090) does not encode the GGI, we applied sequence similarity searches to the GGI of the *Ngo* strain MS11 to identify GGI genes in each isolate. DGI-isolates were more likely to possess the full complement of GGI genes than non-DGI isolates (**Figure 5A**, p-value = 2.96e-05, Welsh’s T-test). A subsequent cluster analysis of the 59 GGI genes across the 301 isolates containing at least one GGI gene revealed a core set of 49 genes present in most (250 of 301) isolates (**Figure 5B**). An additional set of 7 genes are found in 221 of these isolates. Our statistical analyses, however, did not associate any single GGI gene with the DGI phenotype. Combined, this raises the possibility that certain GGI loci might combine with other factors to facilitate a strain’s invasive potential.

**Figure 5:**
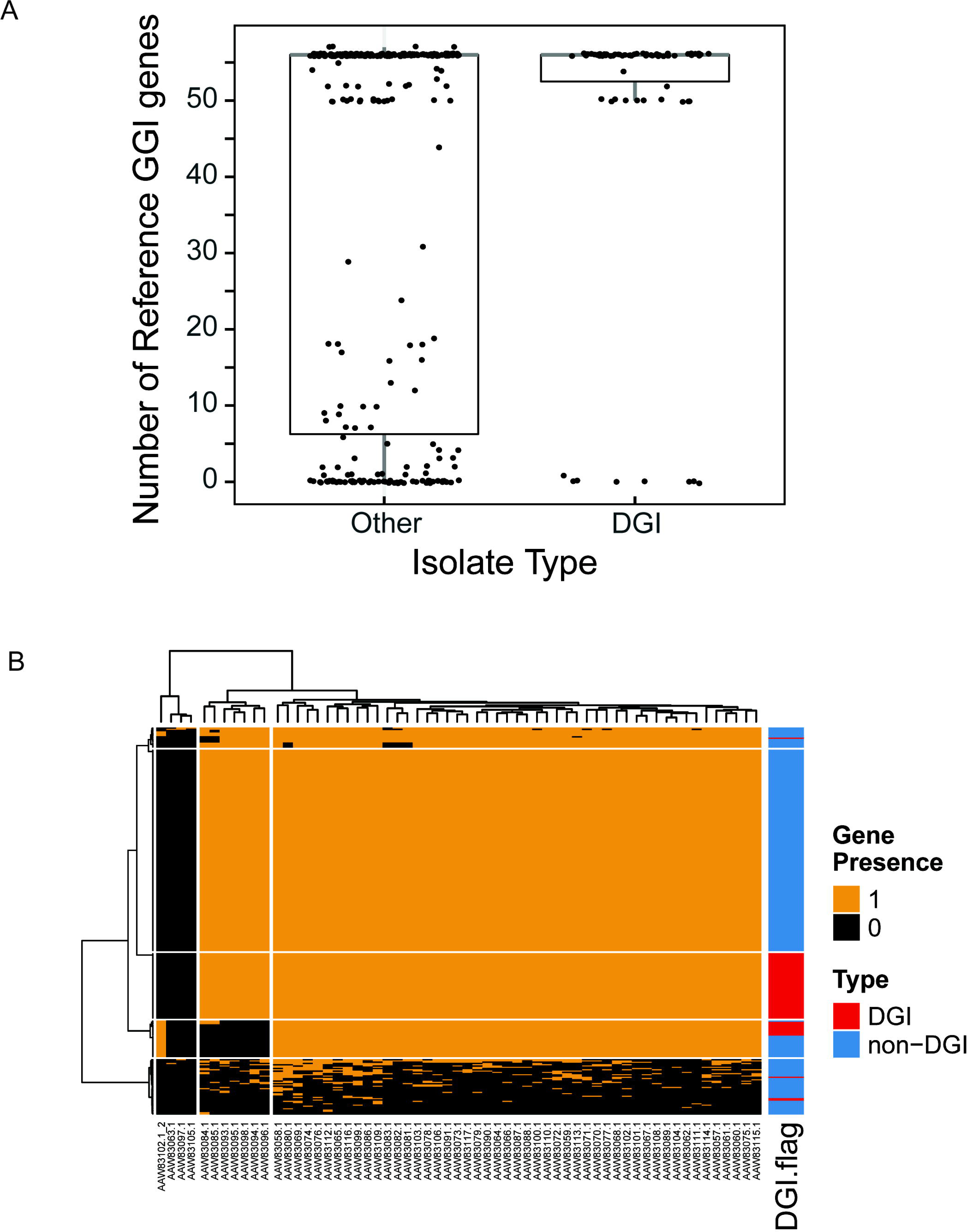
Distribution of GGI-associated genes associated with DGI phenotype. (A) Number of GGI genes in DGI and non-DGI *N. gonorrhoeae* isolates. (B) Presence and absence of 59 gonococcal genetic island genes across *N. gonorrhoeae* isolates.

## DISCUSSION

Amidst the rise of antibiotic resistance and a lack of gonococcal specific vaccines, gonococcal infections represent a pervasive threat to global health. While an infrequent outcome of gonococcal infection, the recent emergence of clusters of DGI is concerning. Invasive gonococcal infections have classically been more frequent in women, presumably because they are more likely to have asymptomatic infections that go untreated, yet the re-emergence of DGI has tended to include men and women in similar proportions [7–10]. Immune suppression is a demonstrated risk factor [25, 26], yet this seems to not be a factor in most recent cases. While a reduced access to healthcare during the COVID-19 pandemic was suggested as a plausible explanation for the increased incidence in California [9], the other incidences began prior to 2019 [7, 8, 10]. The absence of obvious answers has frequently led to suggestion that the increased frequency in cases stems from the emergence of gonococcal isolates with an increased propensity for dissemination [5, 8–10]. The difficulty in exploring this possibility stems from the routine diagnosis of gonococcal infection by nucleic acid amplification testing (NAAT), both for expediency and because *Ngo* can rarely be cultured from infected synovial fluids during DGI [8, 9, 27]. We took advantage of both the diverse gonococcal strain collection of the National Microbiology Laboratories of Canada and the routine isolation of *Ngo* from clinical cases in Ontario to select representative strains for phenotypic analysis and compare full genome sequences by complementary bioinformatic analyses of isolates from uncomplicated (mucosal) and disseminated gonococcal infections.

Besides the possibility of undiscovered/unmeasured epidemiologic or ecologic characteristics, the potential to cause invasive infection has classically, and intuitively, been linked to gonococcal resistance to complement-dependent killing, which occurs because the bacteria is able to sequester human-derived complement regulatory factors to its surface [12, 28]. As with prior studies, most isolates from localized mucosal infections expressed *porB1b* while 74% of DGI isolates encode a *porB1a* allele of the PorB porin, which has been linked to complement regulatory Factor H and C4BP binding [12]. However, the exceptions in this simple relationship indicates that this alone cannot dictate the invasive phenotype.

To extend upon standard typing schemes, we considered whether binding to one or more complement components were a characteristic of the invasive strains, performing an exhaustive flow cytometry-based analysis of complement factor binding after exposing the bacteria to normal human serum. All isolates showed some association with FH and/or C4BP to varying degrees, yet this binding did not strictly correlate with that of C3 or the membrane attack complex (MAC), nor did it predict resistance to the bactericidal activity of serum. We observed that different isolates aggregate to different extents. While this phenotype did not relate to whether the strains were from invasive infection, we considered that the differential association with single versus aggregated bacteria might explain this difference. Indeed, there were some instances where FH and C4BP binding were markedly different between singlets and aggregates in a suspension of the bacteria, however it is not evident how this could explain the marked serum sensitivity (15G0001 and 15G0003) or resistance (18G0016) of these cultures. We repeated the serum bactericidal assays with *Ngo* grown in the presence or absence of CMP-NANA, which confers serum resistance in some strains by virtue of their ability to sialylate their LOS [12], however this did not reconcile the differences. It is plausible that the standardized assay conditions employed, which used 60% pooled normal human serum, may not allow a complete segregation of serum sensitive and resistant bacteria or that immunoglobulin present in the serum differentially interacted with the different isolates. However, it is also likely that the propensity to cause DGI is also facilitated by some other, as yet unrecognized, feature of the bacteria.

The availability of genome sequences generated from hundreds of isolates captured as part of enhanced public health surveillance activities provided us with an opportunity to investigate the genetic determinants related to invasive infection. For this purpose, we included 360 isolates collected across a seven-year time frame (2013-2019) and which was comprised of a broad spectrum of serotypes. When using a traditional NG-MAST analysis, 5 *porB* alleles were prevalent in the Ngo population. Of these, only *2206* and *5136* were significantly associated with DGI. The *2206* allele was spread across all included provinces, with 56 from Manitoba, 27 from Ontario and 7 from Saskatchewan or Alberta, and has been prevalent since 2014. The proportion of DGI isolates from Manitoba and Ontario were comparable. Interestingly, all of the *5136* isolates were from Ontario and seem to have emerged after 2018, and the non-invasive isolates carrying this allele were derived from the eye (**Supp. Table 1**). This makes it plausible to consider that a non-invasive *5136*-expressing isolate obtained some attribute(s) from the invasive *2206* lineage by horizontal genetic transfer.

In considering the remarkable genetic plasticity of *Ngo*, even as a single strain is passaged in culture [29], we applied the population genomic tool, PopNet, to define population structure and identify loci exhibiting shared ancestry across isolates. Applying this tool revealed nine sub-populations, each representing a distinct lineage. This largely separated isolates recovered from Manitoba from those obtained from Ontario, a segregation that was not evident with the SNP-based analysis. Moreover, the most recent Manitoba and Ontario DGI isolates (2018-2019) tended to be clustered together within sub-population A. The fact that certain sub-populations contain no DGI isolates suggests that the potential to cause invasive disease is not uniform among strains. However, with DGI isolates present among 5 of the 9 sub-populations, this reinforces the possibility that the natural genetic competency of *Ngo* has allowed some DGI-associated genetic determinant(s) to be readily shared through genetic exchange.

With our POP-NET-based genomic analyses, we identified 2,657 non-synonymous genomic variants across 975 genes that were enriched in the DGI isolates, making them candidates for a role in DGI. The application of this approach to a broader set of DGI and non-invasive isolates will presumably reduce the number of genes indicated. The lack of strict correlation between any one allele and invasive disease is consistent with the phenotype being multifactorial, and may be further complicated by the fact that mucosal isolates from uncomplicated infection may have the potential to cause DGI. Regardless, certain variants were highly significantly enriched in DGI isolates. These included three different non-synonymous mutations associated with the *purH* gene, which is involved in purine biosynthesis. The functional consequence of these changes is not evident based upon the variant sequences, yet *purH* and other genes upstream in the purine biosynthesis cascade were revealed by a genome-wide mutational screen to facilitate *Haemophilus influenzae* persistence during lung infection in mice [30], supporting the potential for a direct influence on gonococcal virulence. Also provocative, thirteen of the DGI-enriched variants were associated with the cytosolic alginate O-acetyltransferase (NGO_0534, PacA) and four were within the associated periplasmic-localized PacB (NGO_0533), which modify peptidoglycan to confer resistance against hydrolysis by host lysozymes [31, 32]. Classical studies have demonstrated that O-acetylated peptidoglycan from *Ngo* elicits a persistent arthritic response in rats, an effect substantially less apparent with non-acetylated peptidoglycan [33]; this appears to relate to the persistence of undegraded acetylated fragments [34]. Other significantly associated variants occurred in gene products involved in processes including cell envelope integrity, metabolism and pilus assembly, however 7 of the 25 most significant sequences encode hypothetical proteins (**Supp. Table 2**).

Neisserial species can readily transform DNA, and do so to such a degree that the population structure is considered panmictic [35]. Consequently, genetic determinants are readily acquired, including those that may contribute to antimicrobial resistance or pathogenesis [36, 37]. This makes it feasible that the isolates that are causing the recent cluster of disseminated infections have acquired one or more genes not present in the FA1090 reference genome used to define our genetic variants, and would not then be evident from an alignment-based analyses. Indeed, we detected 63 genes that were enriched in DGI isolates but absent in FA1090 (**Table 3**). Six of these, including the two with highest odds ratio, were proteins of unknown function, making them enticing to consider in future studies. Interestingly, four of the DGI-associated proteins are from toxin-antitoxin systems, which are often involved in the control of programmed cell death, stress tolerance or plasmid maintenance [38]. While *Ngo* FA1090 is predicted to encode two toxin-antitoxin systems (*vapBC* and *hicAB*), neither these nor those that we have found to be enriched in the DGI isolates (*prlF*, *ydaS*, *zetA* and *parE*) have previously been linked to pathogenesis. The gonococcal genetic island (GGI), a 57-kb genomic region with a relatively low G+C content, has previously been suggested to carry certain elements that are enriched in isolates from invasive infection [39]. While we did find that the GGI was present in nearly all invasive isolates, we did not identify any individual gene that were significantly enriched in DGI isolates from those that have not been associated with invasive disease.

In this study, we took advantage of the routine collection of gonococcal isolates from both uncomplicated and invasive infections in Canada to allow phenotype analysis focused on bacterial aggregation, serum complement factor binding and resistance to the bactericidal activity of serum, characteristics with clear potential to influence a disseminated outcome. While all isolates possessed the capacity to bind one or both of the complement regulatory proteins C4BP and FH at some level, the specific pattern of binding did not invariably explain the bacterial association with C3 or MAC, and these together did not distinguish disseminated from non-disseminated *Ngo* isolates. A SNP-based genotype analysis of aligned sequences revealed no relationship between DGI isolates since these were scattered among those from uncomplicated disease across the phylogenetic tree. However, a PopNet-based analysis did differentiate Manitoba and Ontario DGI isolates and revealed that their association with invasive disease was not uniform since no cases were apparent within 4 of the 9 subpopulations. This suggests that some feature of the bacteria must influence their capacity to disseminate, however it is not clear if this is inherent in these populations or has been horizontally transmitted from an invasive strain that has previously emerged. Based upon these analyses, it is enticing that there are a variety of non-synonymous genetic differences associated with the DGI phenotype, some of which were shared with the FA1090 but other which were shared among the invasive low passage isolates but absent in this prototype strain. Future studies must consider the potential for these to contribute to increased dissemination or, perhaps counterintuitively, reduced clinical manifestations during mucosal infection so as to promote the incidence of DGI by virtue of more infections being left untreated.

## MATERIAL AND METHODS

### Source of genome sequences and identification of genomic variants

Whole genome sequence data of 360 *Ngo* isolates (available sequences of strains from years 2013-19) were acquired from Public Health Ontario (PHO; Ontario, Canada) and the National Microbiology Laboratory (NML; Manitoba, Canada; **Supplemental Table 1**). Of these, 229 isolates are available at the sequence read archive of the National Center for Biotechnology Information at BioProject IDs PRJNA516461 (27 isolates) and PRJNA533242 (202 isolates). The remaining 131 isolates were acquired from PHO and NML directly.

To identify genetic variants (SNPs, insertions, and deletions), low quality reads were first filtered and trimmed using fastp (v.0.20.1; [40]) with the non-default parameters: qualified_quality_phred = 15; unqualified_percent_limit = 40; cut_right_window_size = 5; cut_right_mean_qu80ality = 20. Reads were aligned to a reference *Ngo* genome (FA1090 strain, genbank ID: NC_002946.2) using BWA (v 0.7.17; [41] with default parameters. Genetic variants were identified using the Genome Analysis Toolkit (GATK) (v.4.1.2.0; [42]). In the absence of a set of high-confidence genetic variants for *Ngo*, base quality and variant quality score recalibrations were not performed. Resulting variants were filtered using the following parameters: QualityDepth < 10.0, FisherStrand > 10.0, StrandOddsRatio > 3.0, RMSMappingQuality < 50.0, MappingQualityRankSumTest < -5.0. Significantly enriched non-synonymous genomic variants in DGI isolates were identified using Fisher’s exact test, with p-values adjusted using Benjamini and Hochberg’s correction with a significance threshold of < 0.20. Functional impact of variants was estimated using SIFT 4G [43].

To determine if mutations across related genes were associated with DGI, a Gene Ontology [44] enrichment analysis of non-synonymous variants was performed. For this analysis, the GO categories of each gene of *Ngo* FA1090 was retrieved from the gene annotations on the UniProtKB database [45]. Each non-synonymous genomic variant was then assigned to its respective GO terms [46]. Variants were ranked according to their p-values from the Fisher’s Test described above, and this list was used for the enrichment analysis, which was performed using the R-package clusterProfiler (v.4.0.5; [47]). Here, p-values of enriched GO-terms were adjusted using the Benjamini and Hochberg (BH) procedure with a significance threshold of 0.05.

### Population analysis

Phylogenetic tree reconstructions were generated from concatenated alignments of single nucleotide polymorphisms (SNPs) using the R-packages phangorn (v.2.11.1; [48]) and ape (v.5.6; [49]). Trees were constructed with the maximum-likelihood algorithm. The optimal model was determined using phangorn’s modelTest function, which resulted in the General Time-Reversible model [50]. Bootstrapping was performed using 100 iterations. Trees were plotted using ggtree (v.3.0.4; [51]).

A population network of the isolates was constructed using the same alignment of SNPs as input into the population analysis tool, PopNet [18]. In brief, using segment lengths of 5000 base pairs, individual sections of each isolate were clustered using the Markov clustering (MCL) algorithm (inflation parameter = 8; pre-inflation parameter = 19). A ‘global’ similarity matrix is then constructed based on the number of segments placed in the same cluster for each pair of isolates, which was clustered again using MCL (inflation parameter = 6; pre-inflation parameter = 2) to define global populations. Genome segments for each isolate are colored based on the population with which it shares most similarity to (chromosome painting). Networks were visualized using Cytoscape (v. 3.8.2) [52]. The similarity of each isolate to one another (defined as the ratio of shared genomic sections) were used to define edges of the graph. In order to reduce the amount of edges in the network and resolve clusters, edges representing under 15% similarity between isolates were removed.

### Identification of non-FA1090 genes associated with DGI

We searched for non-FA1090 genes using the remaining sequence reads that did not align to the reference strain as part of the variant analysis pipeline. For each isolate, the non-aligned reads were assembled *de novo* into contigs using SPAdes (v.3.15; [20]), and putative genes on these contigs were identified using GeneMark (v.1.14_1.25_lic; [21]). Contigs with less than 2X coverage were removed, and putative genes were annotated using the eggNOG mapper [53] against the eggNOG 5.0 database. The ortholog, as defined by the eggNog mapper, was used to annotate the genes [54]. Where unaligned reads assembled into FA1090 reference genes, the annotated genes of FA1090 were annotated using the eggNog mapper and used to filter putative genes matching these annotations. Genes corresponding to genes from 5 neisserial plasmid sequences (Neisserial cryptic plasmid, (NCBI ID: NC_001377; [55]); the prototypical B-lactamase plasmid (NCBI ID: NC_002098; [56]); and three others: NCBI IDs: GU479464.1, GU479465, GU479466; [57]) were similarly annotated. These “non-FA1090” genes were tested for enrichment in DGI isolates using Fisher’s exact test (p<0.2) with BH adjustment for multiple testing.

To identify genes associated with the gonococcal genetic island (GGI), contigs were first aligned to genes associated with the GGI hosted by *Ngo* strain MS11 (NCBI ID: AY803022.1) using BLAST+ (v.2.6.0; [58]); any hits were marked as putative GGI contigs. After this, putative genes on GGI contigs were found using GeneMark. The identity of the putative genes were determined by aligning them against *Ngo* prototype strain MS11 GGI genes using BLAST. The top alignment (by bitscore) determined the identity of the putative gene. A putative gene required a hit with over 80% sequence similarity to be annotated. Differences in the number of GGI genes per isolate between DGI isolates and non-DGI isolates was tested using Welch’s T-test. The presence or absence of each GGI gene per isolate was plotted in a heatmap using ComplexHeatmap [59]. Individual GGI genes were also tested for enrichment in DGI strains using Fisher’s exact test with a significance threshold of 0.20 and using the BH adjustment for multiple testing correction. Only genes that were detected in at least five isolates were used for enrichment testing.

### NG-MAST typing or porB alleles and structural comparison

Sequences assembled de novo with SPAdes (v.3.15) were uploaded to pubMLST for automated NG-MAST typing [60]. Sequences for *porB* alleles of interest were obtained from the pubMLST database and AlphaFold (v2.3.2) was used to predict protein structure. The Protein Data Bank (PDB) Pairwise Structure Alignment tool [61]was used to align structures and determine structural differences between alleles. Allele sequences were aligned using ClustalW [62] and visualized with the R package msa v1.37.3 using TEXshade implemented in the “msaprettyprint” function[63].

### Serum resistance of DGI and n-DGI isolates

*Ngo* cultures were grown overnight on GC agar supplemented with 1% isoVitalex, harvested and resuspended in BHI + 1% isoVitalex (media). OD_600_ was adjusted photometrically to 0.4 (approx. 10^8^ CFU/ml). 40 μl bacterial suspension was mixed with 60 μl of media (M), or human serum (HS), or heat-inactivated human serum (HI). To examine the effect of sialylation on serum resistance, prepared cultures were pre-incubated or not with final concentration of 20 μM CMP-*N*-acetylneuraminic acid (CMP-NANA) at 37°C for 30 mins prior to treating with HS.

### Flow cytometry to assess complement deposition on *Ngo* strains

*Ngo* cultures were grown overnight on GC agar supplemented with 1% isoVitalex, harvested and resuspended in BHI + 1% isoVitalex (media). OD_600_ was adjusted photometrically to 0.4 (approx. 10^8^ CFU/ml). 40 μl bacterial suspension was mixed with 60 μl of media, or human serum, or heat-inactivated human serum. Samples were incubated at 37°C, 5% CO_2_ for either 15 (C3 and C5B-9 factors) or 30 minutes (Factor H and C4BP). Complement activation was stopped by adding 150 μl ice-cold phosphate-buffered saline (PBS)/1% BSA/10 mM EGTA/10 mM MgCl_2_. Samples were processed on ice, and pelleted at 500g, 4°C for 5 minutes and washed twice with ice-cold 0.1% BSA/PBS. The pellet was resuspended and stained with a primary antibody for 45 minutes, washed twice with ice-cold 0,1% BSA/PBS and stained with a secondary antibody for 45 minutes. The samples were washed twice as described above and fixed for 30 minutes with 200 μl of 4% paraformaldehyde solution in PBS. Samples were analyzed by flow cytometry (BD LSRII flow cytometer, Flow Cytometry Facility, UofT) on the same day. Data were analyzed and plotted using FlowJo v10 [64].

Primary and secondary antibodies for different complement factors were used as described: C3-fragments were stained using primary poly-clonal goat-anti-C3 antibody at 1:1000 dilution and secondary donkey anti-goat IgG-Alexa 488 conjugate at 1:500 dilution. C5b-9 was stained using primary mouse anti-human-C5b-9 antibody clone ae11 at 1:1200 dilution and secondary goat anti-mouse Alexa 647 conjugate at 1:500 dilution. Factor H (FH) was stained using primary goat anti-human FH antibody at 1:1000 dilution and secondary donkey anti-goat Alexa 488 conjugate at 1:500 dilution. C4BP was stained using primary rabbit anti-human C4BP antibody at 1:500 dilution and goat anti-rabbit Alexa 647 conjugate at 1:500 dilution.

### Analysis of flow cytometry data

To analyze bacterial populations in FlowJo v10, the main population in a FSC-A/SSC-A scatter plot was selected to exclude cellular debris with low FSC-A or SSC-A. Next, the main population in SSC-W/SSC-A was selected excluding scattering data at the upper and lower limits. In the next step of analysis two distinct populations of aggregates and singlets were defined in FSCH/FSC-A. The singlet population gate boundaries were based on population patterns observed in strains that showed no tendency of aggregation in fluorescence microscopy and transmission electron microscopy. The aggregate population gate boundaries were drawn around the cell populations larger than the singlet population. The same gates were used for all strains in all experiments.

Samples were analyzed for populational parameters regarding the percentage of singlets and aggregates under different experimental conditions, and the quantitative complement deposition in terms of fluorescence intensity. Histograms for aggregates and singlets were overlayed for the final representation using layout editor with Y axis manually adjusted to a maximum value of 500. Geometric mean fluorescence intensity (GMFI) was used to account for scattering data when quantifying complement deposition on samples. Due to strain-dependent variability, GMFI values were normalized to the negative control for each strain to allow better comparison of shifts between strains. For C3 and C5b-9 deposition, samples with heat-inactivated serum were selected as the negative control, since heat-inactivation abolishes complement activity of these factors. For FH and C4BP, samples in media, where these factors are absent, were used as the negative control since these complement factors are heat-stable and active in both heat-inactivated and fresh serum.

### Scanning electron microscopy of Ngo isolates

Bacteria were streaked overnight on GC-agar supplemented with 1% IsoVitalex and incubated at 37°C for 16h. Bacteria was collected using cotton swabs and resuspended in PBS with Ca^2+^ and Mg^2+^, centrifuged at 3000 rpm for 5 minutes twice, and then resuspended in BHI supplemented with 1% IsoVitalex to an OD_600_ approximately 0.1 and grown in shaking incubator at 37°C until the OD_600_ reached 0.4 (typically 4-6 h). Bacteria were collected by centrifugation at 3000 rpm and resuspended in 2 ml phosphate buffer and fixed in equal volume of 3.7% formaldehyde overnight. The samples were then sent for further processing to the Microscopy Facility at the Faculty of Dentistry, University of Toronto. Samples were dehydrated through a series of ethanol rinses (30, 50, 70, 95 and 100%) and critical-point dried with liquid CO_2_. 3000x and 5000x images were processed for each strain sent for analysis.

## ACKNOWLEDGMENTS

(None)

