## Supplementary material for "Phenotypic diversity and shared genomic determinants among isolates causing a large incidence of disseminated gonococcal infections in Canada": Supplementary Figures.pdf

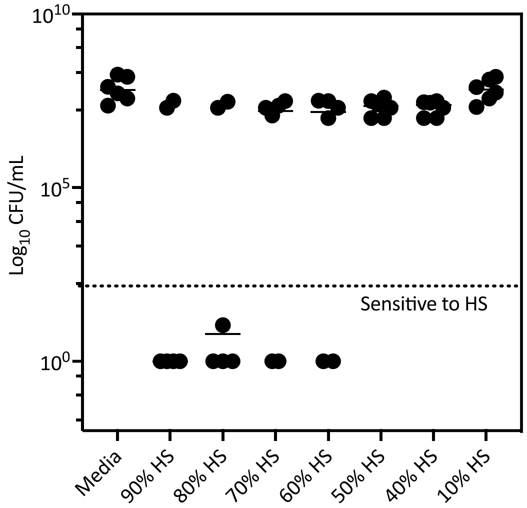


**S. Fig. 1**: Ngo isolates were randomly selected, and bacterial suspension was prepared in PBS with $Mg^{2+}$ and $Ca^{2+}$ to match the absorbance at 600nm~ 0.4, pelleted at 3000rpm and the pellets were resuspended in treatment media as described: media (BHI with 1% IsoVitalex), different concentrations of human serum (10-90%) in BHI with 1% IsoVitalex. Cultures were incubated for 60 min, washed once in PBS with $Mg^{2+}$ and $Ca^{2+}$, serially diluted and plated on GC agar supplemented with IsoVitalex overnight to enumerate for CFUs.

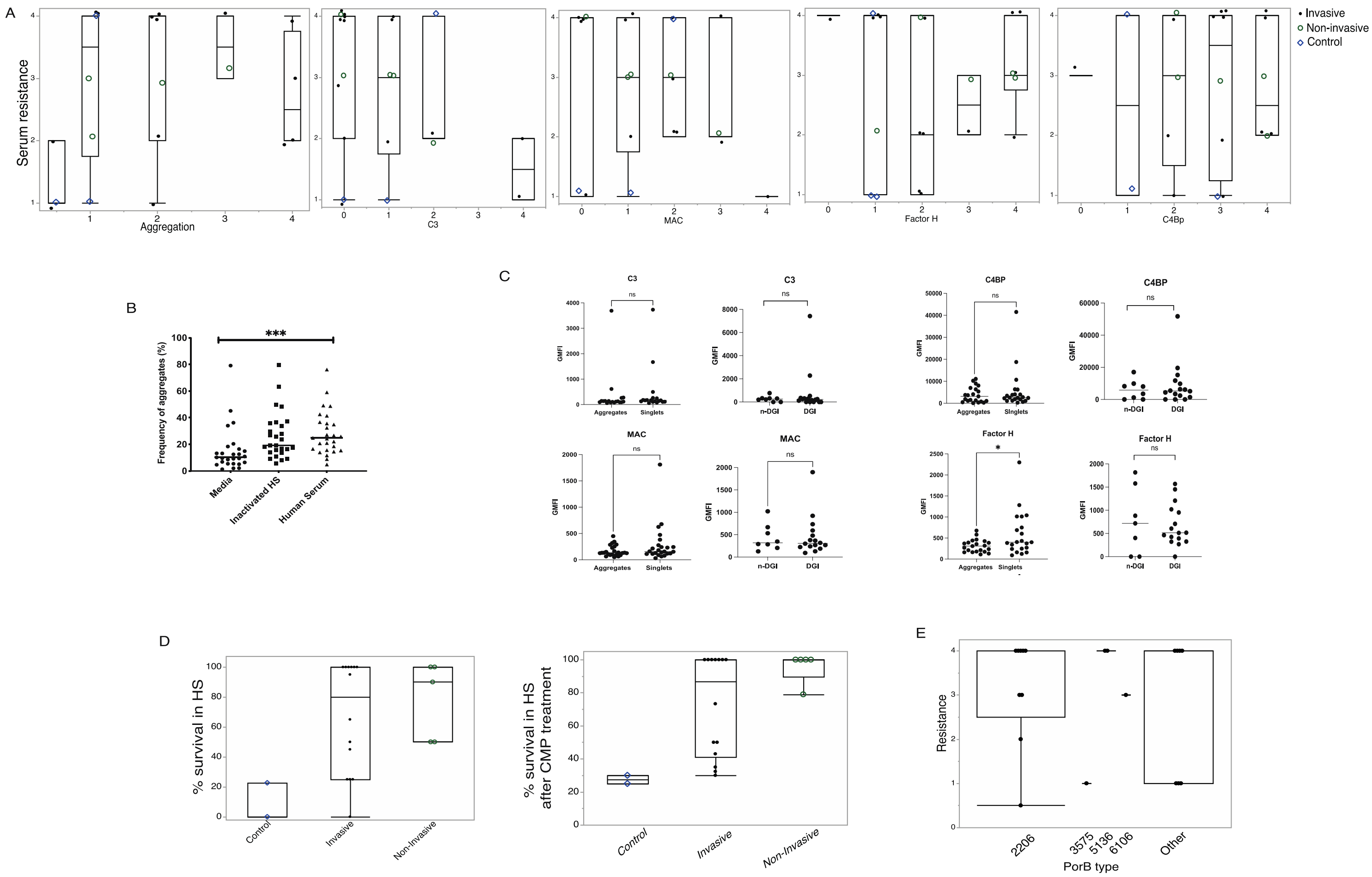


**S. Fig. 2 :** A) Correlation plot for serum resistance to aggregation, C3 deposition, MAC formation Fh and C4Bp recruitment. B) Effect of HS or heat inactived HS on aggregation. C) Comparison between GMFI of various complement factors based on aggregation phenotype or infection type. D) Percentage HS resistance of strains pre-treated with/without CMP-NANA as measured by CFUs. E) Effect of porB allele types on resistance of *Ngo* subset strains against active human serum.

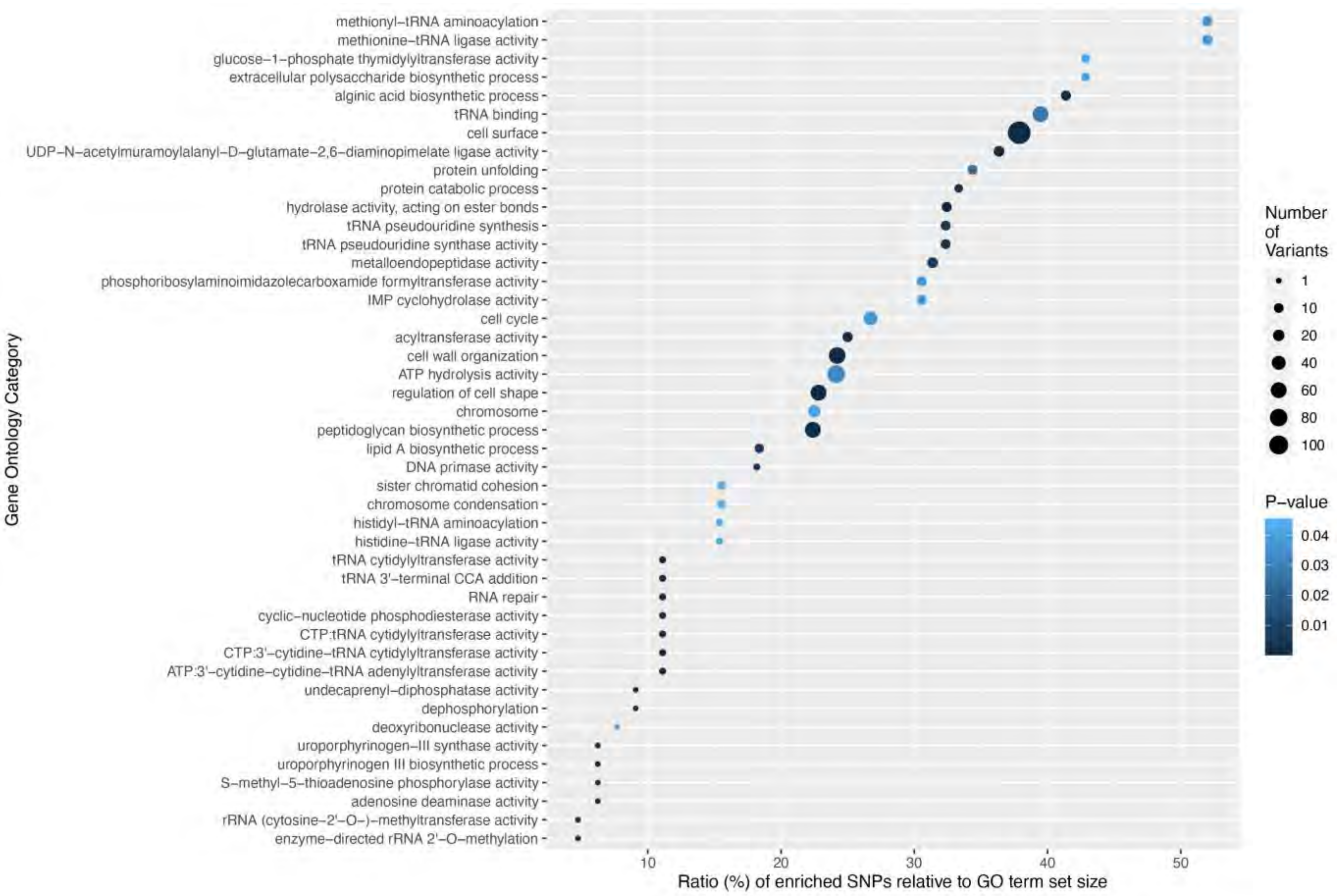


**S. Fig. 3** Functional enrichment of SNPs relative to GO term set size. Size of the circle represents number of variants in each category and the color represents significance (p-value).
